# Population structure and gene flow in the endangered Caribbean reef-building coral, *Acropora palmata*

**DOI:** 10.64898/2026.04.15.718759

**Authors:** Iliana B Baums, Nicolas S Locatelli, Kim L de Luca, Sheila A Kitchen

## Abstract

Populations of the Caribbean reef-building coral, *Acropora palmata*, have declined sharply since their population genetic structure was first characterized in the early 2000s. Previous analyses comprehensively sampled coral colonies across the Caribbean and western North Atlantic but genomic resolution was limited by the number of loci assayed. These analyses indicated extensive asexual reproduction via fragmentation, high outcrossing at the genet level, and a distinct east–west population split. To advance basic research and inform genetic management of this endangered species, we present an updated population genomic assessment using a species-specific microarray to analyze over 4,000 samples representing ∼1,500 genets from 12 geographic regions. Data were contributed by more than 30 research and restoration groups. Our analysis identifies nine spatially structured genetic clusters, with low average pairwise F_ST_ values of between 0.01 to 0.125. Interestingly, legacy genets from the Florida Reef Tract were admixed between two clusters, one dominant in the Mesoamerican Reef Tract on the western flank and the other cluster appearing in genets from Cuba to the south. Migration surface analyses highlight the influence of major current systems on gene flow. Isolation by distance was evident along the Greater Antilles but weak along the Florida Reef Tract. Kinship among wild genets was low across sites, suggesting limited local relatedness; however, assisted sexual reproduction in restoration efforts may disrupt natural kinship patterns. These findings refine our understanding of *A. palmata*’s genetic architecture and underscore the importance of incorporating genomic data into conservation strategies.

## Introduction

Populations of the major reef-building coral species in the Caribbean and northwestern Atlantic have declined dramatically resulting in a significant shift in community composition. Once-dominant major reef-building species including the elkhorn coral *Acropora palmata* and the staghorn coral *A. cervicornis,* along with the massive *Orbicella* species (*O. faveolata*, *O. annularis* and *O. franksi*) are now far less abundant (Gardner et al. 2003). In their place, communities are increasingly dominated by species like *Siderastrea siderea* and *Porites astreoides*, which still display successful natural recruitment but contribute less to reef accretion. These taxa are unable to sustain the high rates of net community calcification necessary for reef framework maintenance (Toth et al. 2023) resulting in a transition from fast-growing, structurally complex *Acropora*-dominated systems to simpler, less resilient reef assemblages (Cramer et al. 2021). Initial declines were driven by eutrophication, sedimentation, disease outbreaks, overfishing, and other direct human impacts (Jackson et al. 2001; Aronson and Precht 2001). More recently, rising sea surface temperatures have intensified coral bleaching and mortality, with global events reaching catastrophic scale in 2023 and 2024 (Reimer et al. 2024).

The Florida Reef Tract has experienced particularly severe losses, especially for *A. palmata*. Genotypic diversity in wild populations has declined steadily since the early 2000s (Baums et al. 2006a). Historical estimates suggest tens to hundreds of thousands of founder genets, but by 2020, only 73 remained in the wild—while approximately 120 were preserved in human care (Williams et al. 2024). As of 2024, *A. palmata* is functionally extinct in the wild along the Florida Reef Tract (Manzello et al. 2025).

The significant losses of both acroporid species in the Florida Reef Tract and the wider Caribbean have been accompanied by an increase in restoration activities (Vardi et al. 2021; Muller et al. 2026). Common strategies include biobanking, fragmenting existing colonies for in situ nurseries, where growth is accelerated before outplanting, and assisted sexual reproduction via gamete collection and cross-fertilization. Gametes are also used to transfer alleles among regions to enhance genetic and phenotypic diversity (Hagedorn et al. 2021; Muller et al.). Such approaches benefit from detailed knowledge of population genetic structure, including genotypic diversity, relatedness, and differentiation, to maximize long-term resilience (Frankham et al. 2017; Baums et al. 2019, 2022).

Prior range-wide studies using five microsatellite markers revealed high genetic diversity, outcrossing, and a distinct east–west population split in the Caribbean with a biophysical barrier near Puerto Rico separating the two regions (Baums et al. 2005, 2006b; Devlin-Durante and Baums 2017). However, limited sampling from the southern Caribbean (e.g., Colombia, Venezuela) hindered precise delineation of the western boundary. Later microsatellite studies confirmed isolation by distance in the Lesser Antilles (Japaud et al. 2019). Additional studies based on single nucleotide polymorphism (SNP) data from *A. palmata* samples that spanned only a limited number of sites from throughout the Caribbean (Devlin-Durante and Baums 2017) and the Mesoamerican Reef Tract (Alvarado-Cerón et al. 2026) have upheld the general pattern of low genetic differentiation among sites within each region.

On a population and reef scale, for any two randomly sampled wild *A. palmata* colonies, there is no detectable level of relatedness (Baums et al. 2006a), indicating low inbreeding risk. However, natural sexual reproduction is now failing in this species, necessitating human-assisted reproduction (Williams et al. 2008; Chamberland et al. 2015). Caribbean acroporids also reproduce extensively via asexual reproduction. Branches break from adult colonies due to physical disturbance and can reattach to the benthos under specific conditions. This can lead to large stands of colonies (ramets) that are nearly genetically identical (clonemates, Baums et al. 2006a). These ramets share the same genome and can trace their origin back to a single sexual reproductive event, forming a genet. In the early 2000’s, the genet-to-ramet ratio in randomly sampled colonies from wild *A. palmata* populations was ∼0.5 (Baums et al. 2006a). *A. palmata* genets can be very old and accumulate somatic mutations in their genome over their lifetimes (Devlin-Durante et al. 2016; Conn et al. 2025). Incidentally, such accumulated mutations can result in negative inbreeding (F) values.

To support conservation and restoration, we present the most comprehensive, range-wide population genomic analysis of *A. palmata* to date, based on 1,502 genets. Data were generated using the species-specific hybridization-based coral-algal genotype array and the Standard Tools for Acroporid Genotyping (STAG, coralsnp.uol.de, Kitchen et al. 2020). Since publication, the array has been used by over 30 groups to process 4,194 *A. palmata* samples (**Table 1** and **Table S1**), of which 4,098 have generously been made publicly available. As the dataset expanded, it revealed an isolated population off Colombia (García-Urueña et al. 2022) - predicted by larval connectivity models (Galindo et al. 2006). Here, we integrate this growing public resource to deliver the most detailed assessment of *A. palmata* population structure in any Atlantic coral species.

**Table 1.** Acropora palmata samples. For each region, the table summarizes the number of unique sites, the number of total samples, the total number of unique genets, the number of unique genets without keywords indicating nursery or assisted reproduction efforts (wild genets), the number of facility-flagged samples (not genets), and the coordinates of the centroid of samples across the region (given in decimal degrees, WGS84 reference frame). Given that not all users distinguished between wild (founder) genets and sexual recruits in the metadata they supplied to the database, the number of “wild” genets may be too high for all regions that engage in conservation and assisted reproduction. Note that the known number of founder genets in Florida is 247 (Williams et al. 2024).

| region | sites | samples | genets | wild_genets | facility_samples | latitude | longitude |
| --- | --- | --- | --- | --- | --- | --- | --- |
| Bahamas | 12 | 60 | 51 | 51 | 0 | 24.0631 | -76.3738 |
| Belize | 31 | 125 | 78 | 71 | 18 | 16.4657 | -88.1990 |
| Colombia | 18 | 80 | 64 | 64 | 0 | 9.8147 | -75.8511 |
| Colombia_SanAndres | 2 | 5 | 5 | 5 | 0 | 14.3625 | -80.1614 |
| Cuba | 3 | 55 | 44 | 44 | 0 | 23.1732 | -82.0993 |
| Curacao | 19 | 196 | 129 | 129 | 0 | 12.0831 | -68.8956 |
| DominicanRepublic | 6 | 89 | 79 | 70 | 11 | 18.4012 | -68.8466 |
| Florida | 199 | 2697 | 651 | 227 | 1813 | 25.0928 | -80.4361 |
| Honduras | 7 | 43 | 35 | 30 | 7 | 16.3000 | -86.5200 |
| Jamaica | 1 | 16 | 15 | 15 | 0 | 18.5228 | -77.7644 |
| Mexico | 8 | 54 | 33 | 26 | 24 | 20.7594 | -86.9500 |
| Mexico_Gulf | 2 | 3 | 3 | 3 | 0 | 22.4770 | -89.7005 |
| PuertoRico | 37 | 477 | 239 | 195 | 91 | 17.9419 | -66.8686 |
| USVI | 15 | 104 | 79 | 79 | 0 | 17.7807 | -64.7163 |
| sum | 360 | 4004 | 1505 | 1009 | 1964 | NaN | NaN |

## Materials and Methods

To assess the presence of population structure across the Caribbean and north-western Atlantic, we utilized microarray data of 4,194 *A. palmata* samples from the publicly available STAGdb database (**Table 1, Table S1**) (Kitchen et al. 2020). The STAGdb database and associated microarray were initially designed using the *Acropora digitifera* reference genome (Shinzato et al. 2011). To convert coordinates to the *Acropora palmata* reference genome (GCF_964030605.1), we mapped the two references against one another using whole-genome alignment with the Progressive Cactus aligner (Armstrong et al. 2020; McKenna et al. 2021). Genomic coordinate systems were adjusted by converting the hierarchical alignment format (HAL) to chain format and using LiftoverVcf in GATK (Auwera and O’Connor 2020) with RECOVER_SWAPPED_REF_ALT=true. From the 53,589 probes provided by STAGDB as a downloadable VCF file (“all_genotypes.vcf”; *dataset 1*), only recommended genotyping probes (Kitchen et al. 2020) were preserved after liftover (n=18,898 SNPs), low quality genotypes (CONF>0.01) and sites with >10% missing data were removed using bcftools (Danecek et al. 2021), and samples with > 5% missing data were removed (*dataset 2*, n=18,258 SNPs, n=1,397 samples). Kinship was estimated using Plink2 (--make-king-table) and close kin (ramets of the same genet and second degree or closer relatives) were removed with a KING kinship threshold (king-cutoff) of 0.08834 in Plink2 (Chang et al. 2015). The pruning strategy implemented in Plink2 maintains only one sample from each family where “family” is defined as clusters of related samples which include ramets of the same genet as well as closely related genets and their offspring. The sample with the least amount of missing data per family was retained. Nine additional samples were excluded due to exceedingly low heterozygosity, indicative of inbreeding or selfing, some of which were generated by assisted reproduction (Vasquez Kuntz et al. 2022). For isolation-by-distance analyses (see below for additional methodological information), we performed minimal additional filtering, keeping biallelic SNPs, removing only globally rare alleles (minor allele count >3), and pruning for linkage disequilibrium (R^2^ < 0.5 in 100Kb windows with bcftools +prune; *dataset 3*, n=5,555 SNPs, n=955 samples). For population structure analyses (PCA, STRUCTURE/ADMIXTURE, F_ST_, F, and heterozygosity), we filtered more stringently, downsampling overrepresented regions such that no geographic region had more than 60 samples represented in downstream population genomic analyses. We subsequently only retained unlinked (R^2^ < 0.5 in 100Kb windows) biallelic SNPs with minor allele frequency >0.05 (*dataset 4*, n=3,215 SNPs, n=554 samples, **Table 1, Table S1**). To illustrate the importance of thorough, microarray-specific probe filtering, we provide a comparison of our most filtered dataset (*dataset 4*), to a dataset where only kinship and MAF>0.05 filtering was performed (*dataset 5*, **Figure S1**), as well as comparison between *dataset 4* and a dataset where all filtering was performed, with the exception of filtering for CONF which is a format field unique to microarray data types (*dataset 6*, **Figure S2**).

With filtered variant sites (dataset 4), STRUCTURE was run to infer population structure using values of K=1 to K=12 (Pritchard et al. 2000). For each value of K, ten replicate chains were run with a burn-in of 50,000 and a further 50,000 steps were preserved. For each run, chains were visually inspected to ensure convergence. All runs were performed using the admixture model without a population prior. ADMIXTURE (Alexander et al. 2009) was also run, using K=1 to K=12, each with ten replicate runs and 10-fold cross validation. StructureSelector (Li and Liu 2018) was used to run CLUMPAK (Kopelman et al. 2015) and identify modes across runs to evaluate the most likely values of K using the Puechmaille (Puechmaille 2016), delta K (Evanno et al. 2005), and cross validation error minimization methods. To visualize effective migration surfaces and barriers to migration, FEEMS was run on a global triangular grid with a resolution of 8 (corresponding with a cell area of 7,774km^2^) and cross validation using the previously filtered dataset across 10 log-evenly spaced intervals from lambda = −2 to lambda = 2 (Marcus et al. 2021). Where colony or reef GPS coordinates were missing, approximate reef coordinates were assigned based on the reef name or region submitted by the user (**Table S1**).

To estimate pairwise F_ST_ between sampling locations using microarray data, we used Weir and Cockerham (1984) estimation with the R package StAMPP (Pembleton et al. 2013) and calculated bootstrapped F_ST_ values from dataset 4 (unlinked biallelic SNPs). With the same unlinked SNPs, we also subset the data to genets from a given region (based on STAGdb region metadata), filtered for MAF >0.05 within the subset, estimated inbreeding coefficients (*F_IS_*) and calculated observed heterozygosity, as implemented in VCFtools (Danecek et al. 2011). Pairwise F_ST_ values were transformed to estimate migration rates under Wright’s island-model approximation N_m_ = (1 - F_ST_) / (4 * F_ST_) where N is the effective population size and m is the migration rate.

Isolation by distance (IBD) was evaluated along the Greater Antilles and along the Florida reef tract by comparing pairwise genetic and geographic distances among sampling sites. IBD analysis was performed on dataset 3, without geographic downsampling but including the same global minor allele call rate > 3 as above and a regional minor allele frequency > 0.05. Wild samples were retained based on metadata by excluding reef names with keywords [“nursery”, “restoration”, “outplant”, “OP”, “situ”, “cross”, “recruit”, “sexual”, “batch”, “MML”, “spawn”, “FLAQ”, “gene”] (**Table 1**). Additional samples were excluded in two further steps: first, by pattern-based reef name matching, removing sites associated with known nursery or restoration facilities (Coral Restoration Foundation, Mote Marine Laboratory, Florida Aquarium); second, by coordinate-based matching, which removed any remaining samples sharing the exact geographic coordinates of a confirmed facility site regardless of its reef name label.

Duplicate or closely related reef labels were consolidated where appropriate (**Table S1**). Nearby sampling locations within each geographic subregion were consolidated using complete-linkage hierarchical clustering: sites within 10 km of each other were merged into a single site, with the canonical name assigned to the member with the greatest genet count. Only sites with >5 unique genets per site were retained for IBD analysis. Genet identifiers were obtained from the STAGdb “export_all_samples_data.tabular”. STAGdb identifies genets based on a threshold of pairwise genetic distancelJ=lJ0.032 and implemented in the multilocus genotyping tools of the STAGdb workflow (Kitchen et al. 2020). Genetic distance was expressed as linearized F_ST_, calculated as F_ST_ / (1 - F_ST_), using the Weir and Cockerham (1984) estimator implemented in Python package scikit-allel (Miles et al. 2024). Geographic distance between sites was measured as great-circle distance using the haversine formula (Greater Antilles). Sites were additionally ordered along the Florida reef tract using a greedy nearest-neighbor approach, followed by calculating cumulative along-tract distances. For the Florida dataset, sites were assigned to Upper, Middle, or Lower Florida Keys sections by nearest-centroid classification using geographic reference points. The association between genetic and geographic distance matrices was evaluated with Mantel tests (Pearson correlation, 10^4^ permutations) implemented with skbio.stats.distance.mantel. Effect size was summarized by fitting an ordinary least squares regression and reporting R^2^ (scipy). Data handling and plotting was done with numpy, pandas, matplotlib, and plotly. Full code is available for metadata curation, sample filtering, and IBD analyses.

## Results

We analyzed publically available genotyping data from 4,194 samples of the endangered reef-building coral, *Acropora palmata*. To satisfy assumptions of population genetic models, it is necessary to filter the genotyping data prior to analysis for genotyping probes and sample quality. The coral-algal microarray was designed with > 53,000 genotyping probes for *A. palmata* as well as *A. cervicornis* and for coral algal symbionts (genera in Symbiodiniaceae), however, only a subset meet the quality thresholds for genotype assignment (Kitchen et al. 2020). Sample data was first filtered to include the previously determined best and recommended probe set for the acroporoids (Kitchen et al. 2020) and then further filtered to retain *A. palmata* population variable probes. Applying common data filtering thresholds on all probes instead of using the best and recommended probe set can obscure sample differentiation (**Figure S1**). Similarly, failure to exclude low quality genotype calls prior to downstream analysis can yield genetic groups that are driven by technical errors (**Figure S2**). An additional critical step for population genetic analysis of asexually reproducing species is the assignment of ramets to genets. Samples were assigned to genets based on previously established, rigorously tested pairwise genetic similarity thresholds that take into account within-genet SNP variation based on biological processes (somatic mutations) and technical error (microarray genotyping, Kitchen et al. 2020). Only unique genets were considered in the subsequent analyses (n = 1,502, **Table 1**).

Population structure and migration analyses using microarray data were conducted on a geographically balanced subsampled dataset of 554 genets (max 60 genets per region). These analyses support previous findings that *A. palmata* genets are separated into distinct populations across the Caribbean (**Figure 1**). The two first axes of the principal component analysis (PCA) explained 10% and 4% of the variation in the data. As expected, the PCA recapitulates the geography of the sampling locations. The first axis separates sampling locations along an east-west axis, with the Mesoamerican reef locations largely overlapping (Mexico, Belize, Honduras). About half of the genets from Florida align with this group. The other half of the genets from Florida group with samples from the Gulf side of Mexico. A handful of the genets from Florida grouped together with samples from Cuba. The samples from the Bahamas and the Dominican Republic formed partially overlapping groups along PC1. Puerto Rico and the US Virgin Islands each formed recognizable groups. Along PC2, samples from the Bahamas, Jamaica and Cuba separated to some degree. Samples from Curaçao formed their own group that was more closely aligned with the Greater Antilles sampling locations than samples from the mainland of Colombia. Indeed, samples from mainland Colombia and the San Andrés Islands formed the most distinctive group in the PCA.

**Figure 1.**
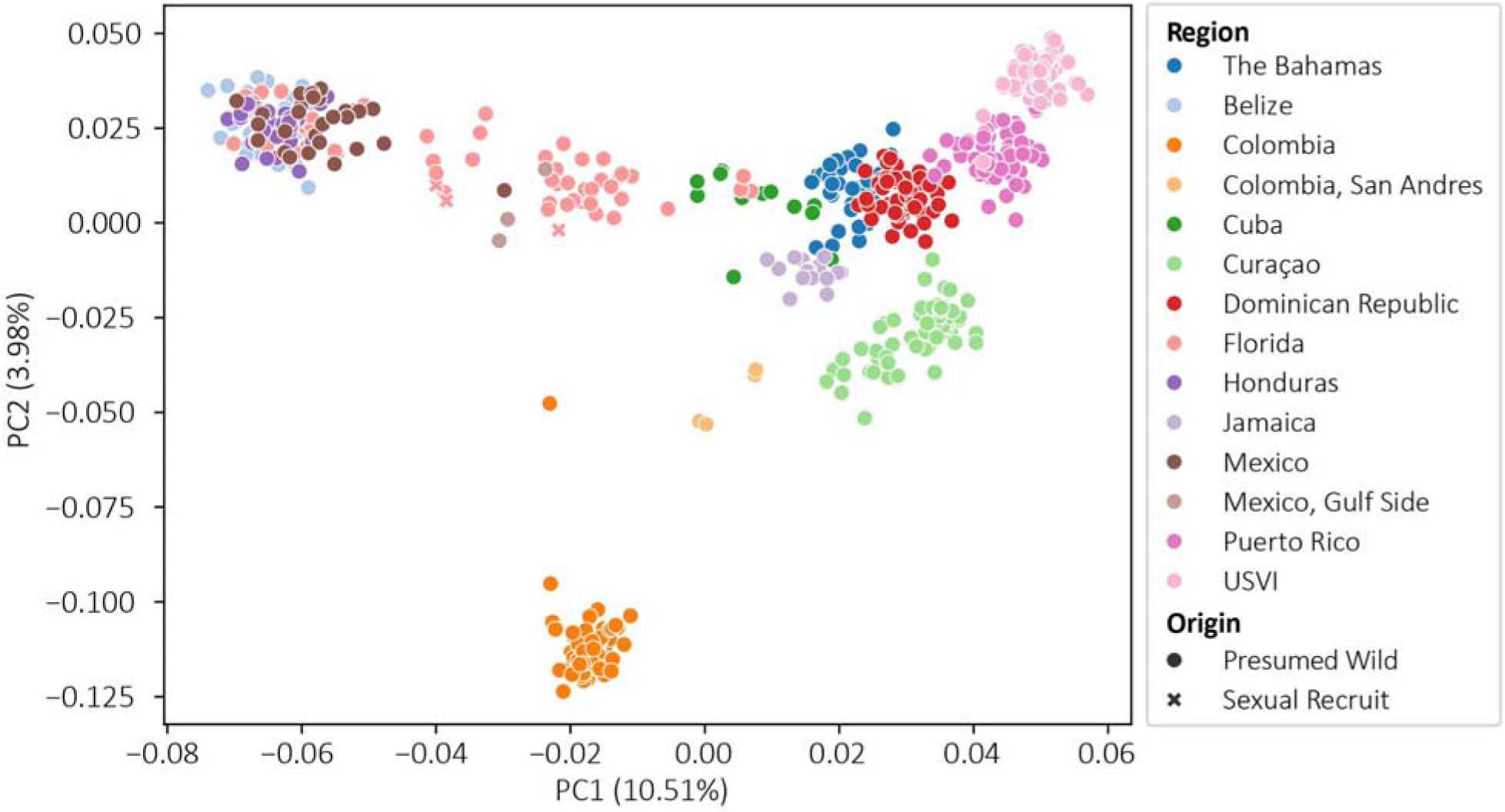
Principal component analysis of 554 *Acropora palmata* genets based on 3,223 SNP loci. A principal component analysis was performed on genotypes assigned to genets via the standardized *Acropora* Galaxy pipeline (Kitchen et al. 2020). The dataset was filtered to remove linked loci, low base calling confidence, site and genet level missingness. The first two principal components are displayed with each point representing a single genet. Shapes denote whether the genet is a founder (circle) or sexual recruit (cross). Colors are assigned to countries where samples were collected: Bahamas = dark blue, Belize = light blue, Colombia = dark orange, San Andrés archipelago = light orange, Cuba = dark green, Curaçao = light green, Dominican Republic = red, Florida =coral, Honduras = purple, Jamaica = light purple, Mexico - Caribbean side = brown, Mexico- Gulf side = tan, Puerto Rico = magenta, US Virgin Islands = light pink.

Unsupervised clustering by STRUCTURE resulted in the division of the 554 genets into K=9 populations, determined to be the most suitable K according to the Puechmaille method (**Figure 2A**). The Evanno method instead supported K=2 as the optimal K (**Figure S3**). Analysis of the same dataset with ADMIXTURE also yielded an optimal K = 9 (according to the Puechemaille method, **Figure S4**) and is therefore considered the best solution. In the western Caribbean, samples from Mexico, Belize, and Honduras clustered together into a Mesoamerican population similar to the results of the PCA. STRUCTURE analysis indicated gene flow into Florida from the Mesoamerican population and between Cuba and Florida. This is evident in the high assignment probabilities (>90%) of some Florida genets to the Mesoamerican population consistent with recent gene flow. The limited number of genets from Cuba (n=13) were admixed themselves with apparent contributions from a cluster dominant in the Bahamas and a second cluster shared with Florida. Dominican Republic genets had varying degrees of ancestry in the Bahamian/Jamaican cluster but assignment probabilities were well below 50% indicating that gene flow among the clusters occurred some generations ago. In the eastern Caribbean, Puerto Rico and the US Virgin Islands each formed a cluster with minor US Virgin Islands ancestry in several Puerto Rican genets. Genets from the US Virgin Islands, Curaçao, and Colombia had high assignment probabilities to their respective clusters, indicating low gene flow from other clusters into these sites.

**Figure 2.**
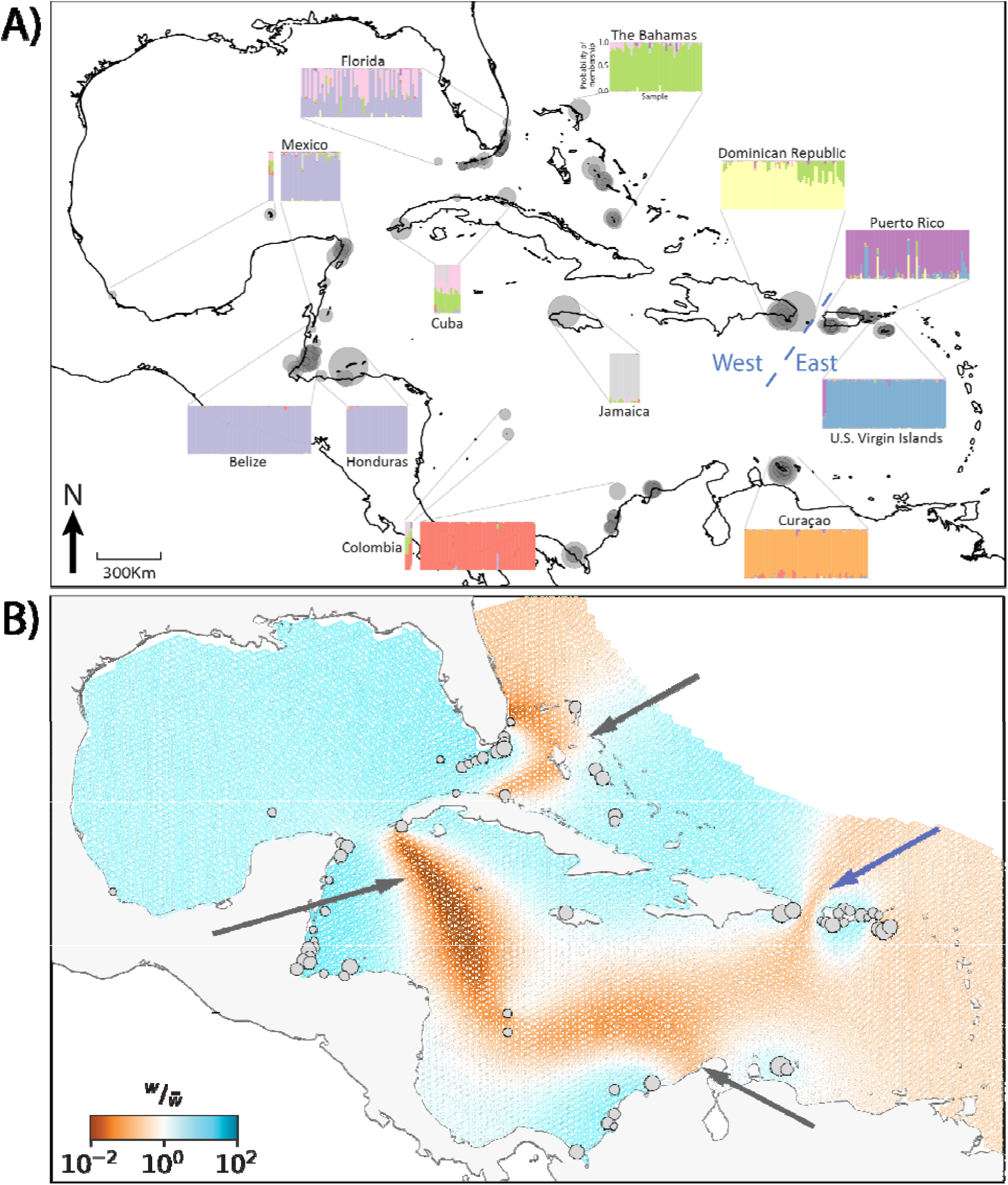
STRUCTURE (A) and effective migration surfaces FEEMS (B) for *Acropora palmata* in the Caribbean. A) The STRUCTURE (Pritchard et al. 2000) results of the unlinked SNPs are presented on a map of the Caribbean and north-western Atlantic with the optimal K value of 9 determined by the Puechmaille method (Puechmaille 2016). For each region, the probability of cluster membership for K = 9 (Y-axis) is plotted for each genet (x-axis). B) Spatial patterns of genetic variation were inferred using FEEMS (Marcus et al. 2021) based on the unlinked SNPs. GPS coordinates were shifted automatically to overlap with nodes on the global grid. Sampling locations are shown as grey circles whose size corresponds with sample size. Color gradients indicate the relative migration rates (W/W), with orange hues corresponding to lower-than-expected migration and blue hues corresponding to higher migration. Dark orange regions may act as barriers that limit gene flow between regions. The blue arrow indicates a previously identified bio-physical barrier to gene flow between the eastern and western Caribbean. Additional barriers identified here are indicated with grey arrows.

While cluster analyses supported the presence of barriers to gene flow throughout the range, genetic differentiation (F_ST_) between sites remains low. For example, samples from neighboring Belize and Honduras belong to the same genetic cluster (population) and have a pairwise F_ST_ of 0.001 (95% CI [0.001, 0.002]; p<0.05). Samples from Jamaica and the Dominican Republic are also geographically close but have a 47 times higher pairwise F_ST_ of 0.041 (95% CI [0.037, 0.045]; p<0.05). The highest pairwise F_ST_ of 0.125 was observed between the distant sites of Belize and the US Virgin Islands (95% CI [0.116, 0.133]; p<0.05). All microarray-derived F_ST_ values and confidence intervals are presented in **Table 2**.

**Table 2.** Pairwise F_ST_ for all regions sampled with confidence intervals (above diagonal) and F_ST_ values (below diagonal). F_ST_ values and confidence intervals were estimated using STAMPP (Pembleton et al. 2013) with 100 bootstraps. All values were significant (p<0.01). FL=Florida, DR=Dominican Republic, JM=Jamaica, MX=Mexico, CU=Curaçao, VI=US Virgin Islands, BE=Belize, PR=Puerto Rico, BH=the Bahamas, HN=Honduras, CB=Cuba, CO=Colombia, GOM=Mexico, Gulf Side, SA=San Andrés Archipelago, Colombia.

|  | BH | BZ | CO | SA | CB | CU | DR | FL | HN | JM | GOM | MX | PR | VI |
| --- | --- | --- | --- | --- | --- | --- | --- | --- | --- | --- | --- | --- | --- | --- |
| BH |  | [0.086, 0.097] | [0.083, 0.096] | [0.027, 0.039] | [0.017, 0.023] | [0.056, 0.064] | [0.022, 0.026] | [0.046, 0.052] | [0.084, 0.096] | [0.029, 0.034] | [0.053, 0.068] | [0.073, 0.083] | [0.034, 0.04] | [0.045, 0.055] |
| BZ | 0.091 |  | [0.084, 0.095] | [0.069, 0.082] | [0.076, 0.086] | [0.105, 0.117] | [0.094, 0.105] | [0.015, 0.018] | [0.0, 0.002] | [0.091, 0.104] | [0.038, 0.05] | [0.002, 0.004] | [0.105, 0.118] | [0.119, 0.133] |
| CO | 0.089 | 0.09 |  | [0.036, 0.047] | [0.082, 0.091] | [0.085, 0.096] | [0.087, 0.099] | [0.065, 0.074] | [0.085, 0.096] | [0.079, 0.089] | [0.076, 0.095] | [0.08, 0.091] | [0.096, 0.108] | [0.113, 0.128] |
| SA | 0.033 | 0.075 | 0.041 |  | [0.023, 0.035] | [0.049, 0.063] | [0.036, 0.049] | [0.03, 0.04] | [0.069, 0.083] | [0.033, 0.046] | [0.037, 0.058] | [0.056, 0.068] | [0.044, 0.056] | [0.061, 0.076] |
| CB | 0.02 | 0.081 | 0.087 | 0.03 |  | [0.064, 0.073] | [0.037, 0.045] | [0.031, 0.036] | [0.076, 0.085] | [0.028, 0.037] | [0.043, 0.057] | [0.063, 0.071] | [0.048, 0.058] | [0.061, 0.073] |
| CU | 0.06 | 0.111 | 0.09 | 0.056 | 0.069 |  | [0.054, 0.063] | [0.071, 0.08] | [0.106, 0.118] | [0.07, 0.079] | [0.075, 0.09] | [0.098, 0.11] | [0.043, 0.05] | [0.056, 0.067] |
| DR | 0.023 | 0.1 | 0.092 | 0.043 | 0.042 | 0.058 |  | [0.057, 0.065] | [0.093, 0.105] | [0.038, 0.045] | [0.067, 0.083] | [0.083, 0.092] | [0.029, 0.034] | [0.045, 0.056] |
| FL | 0.049 | 0.017 | 0.069 | 0.035 | 0.033 | 0.076 | 0.062 |  | [0.015, 0.019] | [0.051, 0.059] | [0.017, 0.026] | [0.012, 0.014] | [0.067, 0.077] | [0.08, 0.091] |
| HN | 0.09 | 0.001 | 0.09 | 0.075 | 0.08 | 0.112 | 0.1 | 0.017 |  | [0.093, 0.105] | [0.037, 0.049] | [0.002, 0.004] | [0.106, 0.119] | [0.118, 0.133] |
| JM | 0.031 | 0.098 | 0.084 | 0.039 | 0.033 | 0.074 | 0.041 | 0.055 | 0.099 |  | [0.073, 0.088] | [0.081, 0.091] | [0.052, 0.06] | [0.068, 0.078] |
| GOM | 0.061 | 0.044 | 0.084 | 0.048 | 0.05 | 0.082 | 0.076 | 0.022 | 0.043 | 0.082 |  | 0.037 | [0.075, 0.091] | [0.088, 0.105] |
| MX | 0.077 | 0.003 | 0.085 | 0.062 | 0.067 | 0.104 | 0.088 | 0.013 | 0.003 | 0.086 | [0.032, 0.042] |  | [0.095, 0.108] | [0.105, 0.121] |
| PR | 0.037 | 0.112 | 0.102 | 0.05 | 0.053 | 0.046 | 0.031 | 0.072 | 0.113 | 0.055 | 0.083 | 0.101 |  | [0.02, 0.028] |
| VI | 0.051 | 0.125 | 0.12 | 0.069 | 0.067 | 0.061 | 0.051 | 0.086 | 0.125 | 0.073 | 0.097 | 0.113 | 0.024 |  |

Effective migration surface analysis served to illustrate the routes of and barriers to *A. palmata* dispersal in the region (**Figure 2B**). The most striking observation is the lack of barriers to gene flow from the Mesoamerican Barrier Reef into Florida. Direct dispersal between Florida and the Bahamas was impeded while dispersal into Florida may occur from Hispaniola and Jamaica via Cuba. As in previous studies, a barrier to gene flow was identified between the Dominican Republic and Puerto Rico. Curaçao and Colombian mainland sites were each isolated.

We found that all regions exhibited similar median genome-wide heterozygosity estimates (39.3-41.6%) with the lowest heterozygosity in Colombia (39.3%) and the highest in Jamaica (41.6%) (**Table 3**). Median inbreeding coefficients (*F_IS_*) were low within each location and ranged from -0.008 in Jamaica to 0.0446 in Florida (**Table 3**). The higher F_is_ value in Florida can be explained by admixture of two clusters across the Florida reef tract (Wahlund effect) or by increased levels of somatic mutations (less likely to be detectable by the microarray).

**Table 3.** Inbreeding and heterozygosity values for *Acropora palmata* for each focal region as determined by microarray SNP data. Inbreeding coefficients (*F*) and microarray SNP heterozygosity (heterozygosity at only variant sites) were calculated using the --het function of VCFtools. Sample sizes were below the minimum recommended for calculating inbreeding coefficients (Gulf Side of Mexico (n=3) and the San Andres Islands of Colombia (n=4), and are indicated with an asterisk (*).

| | Inbreeding $F$ | Observed Heterozygosity |
| --- | --- | --- |
| The Bahamas | 0.0136 | 40.76% |
| Belize | 0.0212 | 39.91% |
| Colombia | 0.0134 | 39.31% |
| Colombia, San Andres | 0.0247* | 44.63%* |
| Cuba | 0.0344 | 41.24% |
| Curaçao | 0.0211 | 40.36% |
| Dominican Republic | 0.0212 | 40.12% |
| Florida | 0.0446 | 40.29% |
| Honduras | 0.0102 | 40.35% |
| Jamaica | -0.0083 | 41.61% |
| Mexico, Gulf Side | 0.0214* | 47.41%* |
| Mexico | 0.0194 | 40.51% |
| Puerto Rico | 0.0195 | 40.63% |
| US Virgin Islands | 0.0097 | 40.64% |

Splitting Florida samples by STRUCTURE cluster (>50% ancestry to one cluster) yielded slightly lower F_is_ (0.032 and 0.024 for the Cuba and Mesoamerican cluster of Florida, respectively) and similar heterozygosity values to those shown in **Table 3**. Pairwise KING kinship values were centered around 0 (unrelated) across all sites where there were >10 genets with which to calculate pairwise kinship coefficients (**Figure 4**). With the exception of two sites in the Dominican Republic, most putatively wild and unaltered *A. palmata* sites did not harbor genets that were closely related to one another. However, sites with known restoration activity such as Sea Aquarium in Curaçao exhibit longer tails and multimodality, indicating higher than expected kinship due to human interventions **(Figure 4)**

Isolation-by-distance analyses revealed significant relationships between geographic distance (expressed either as haversine distance or along-tract distance) and genetic distance (expressed as linearized F_ST_) along the Greater Antilles (Dominican Republic, Puerto Rico, US Virgin Islands; Mantel test r = 0.9; p < 0.01, n = 91) but not the Florida Reef Tract (Mantel test r = 0.2; p > 0.05, n = 45) (**Figure 3**). Only sites with more than 5 unique genets were included in this analysis, but sampling effort was not standardized when samples were collected, limiting the robustness of the results.

**Figure 3.**
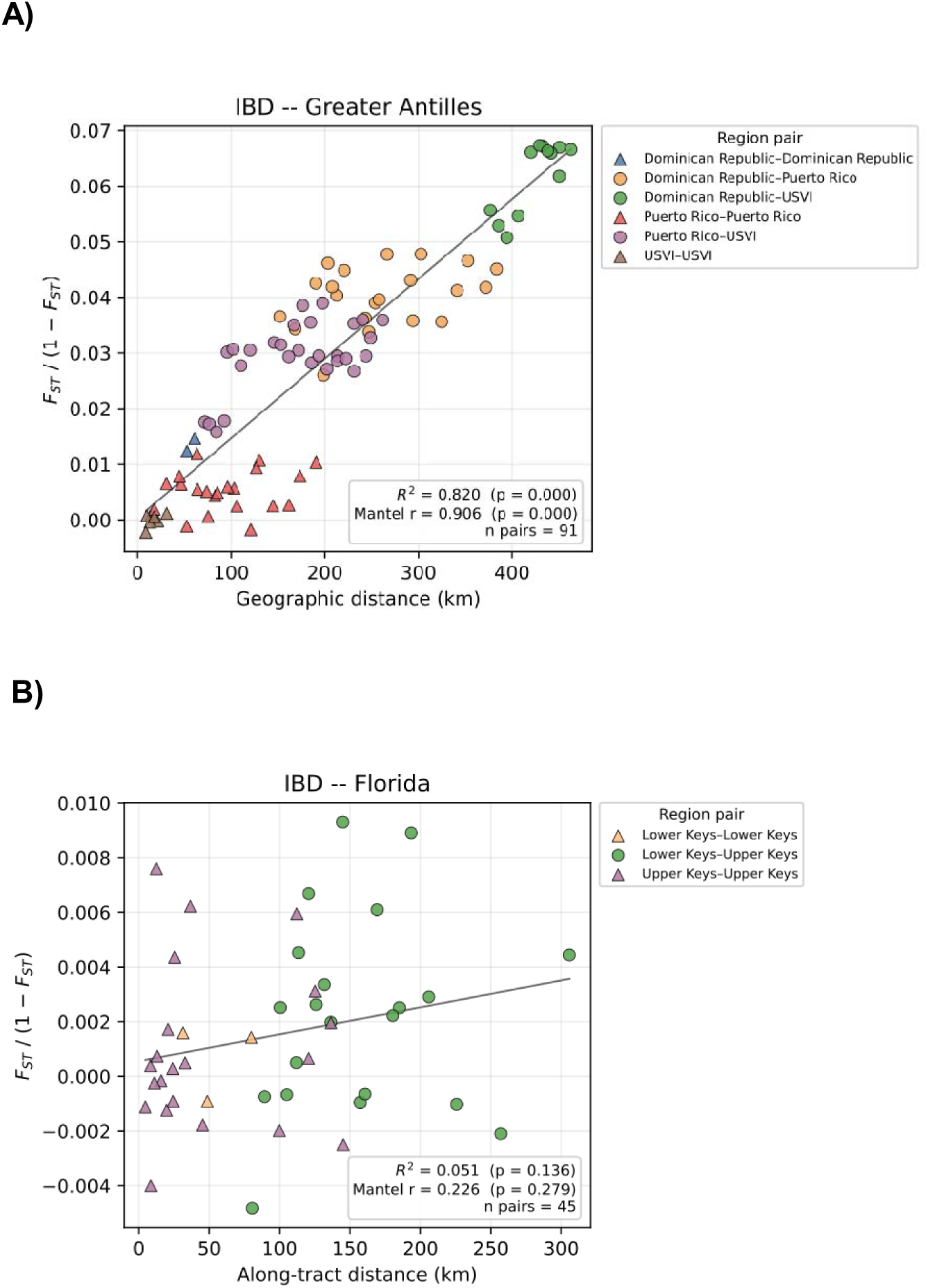
Isolation-by-distance analysis for two regions in the Caribbean and north-western Atlantic. Panels include A) comparisons over 400 km along the Greater Antilles including sites from the Dominican Republic, Puerto Rico and the US Virgin Islands; and B) comparisons over 250 km along the Florida Reef Tract. Plotted are the pairwise geographic haversine (A) or along-tract (B) distances (km) between sites and the linearized pairwise F_ST_ values (unitless) between sites, colored by comparison (within and between regions). Only sites with more than 5 unique genets were included and sites <10 km apart were merged. Results of an ordinary least squares regression (OLS) and a Mantel test of association between geographic and genetic distance are given.

**Figure 4.**
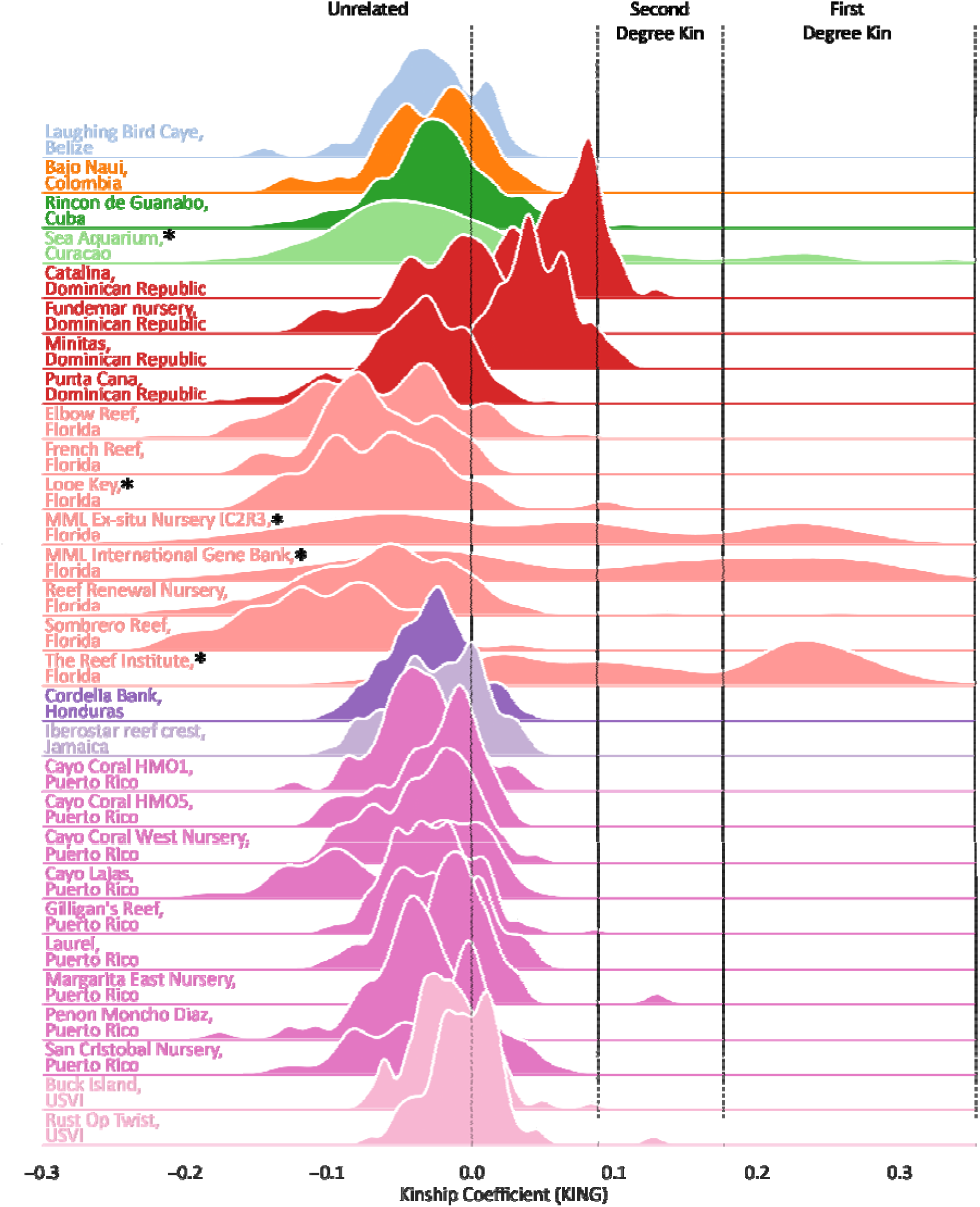
**Within-site relatedness of A. palmata genets**. Genets from wild *A. palmata* stands are generally unrelated to each other while assisted sexual reproduction can increase relatedness among genets at a site or in a biobanking facility. Shown are density distributions of the KING robust kinship values by site, colored by region for sites with >10 genets. Sites and biobanks known to hold genets that resulted from assisted sexual reproduction are indicated with a black asterisk (*). Note that not all sexual offspring are flagged in the STAGdb user-supplied metadata.

## Discussion

The iconic elkhorn coral, *Acropora palmata,* is one of only five reef-building species in the Caribbean and north-west Atlantic. Populations have declined drastically since the first region-wide assessment of population genetic structure was published in 2005. Here, we provide a comprehensive update based on 3,223 genome-wide SNP loci assayed using a high-fidelity microarray.

Unsupervised clustering of genotypes from a subsampled dataset of 554 genets collected across 12 regions and approximately 223 user-submitted sites/reefs resolved into 9 genetic clusters (**Figures 1 & 2**). Clusters largely correspond to geographic regions, with notable exceptions. Genets along the Florida Reef Tract exhibit admixed ancestry, reflecting contributions from a cluster found in Florida and Cuba and a cluster found along the Mesoamerican Reef Tract. Admixture patterns are common in coral genetic data, including in the sister species *A. cervicornis* along Florida’s reefs (Drury et al. 2016; Torquato et al. 2022). The admixed ancestry present in Florida does not appear to be a recent phenomenon as the samples included in the analysis include later generation backcrosses between the Mesoamerican/Floridian and Cuban/Floridian genetic clusters. STRUCTURE analysis did not identify a cluster that contained genets with high assignment probabilities (>75%) to the Cuba/Florida cluster which may be found in an as yet unsampled locality or purebred genets of this cluster may be extinct. While the origin of the Florida/Cuba cluster remains a mystery, biophysical modeling supports the conclusion that the Mesoamerican Reef Tract serves as the origin of the Mesoamerican/Floridian cluster because dispersal is strongly unidirectional from the Mesoamerican Reef Tract to Florida (Baums et al. 2006b; Galindo et al. 2006). These modeled dispersal pathways are corroborated by migration surface analyses in FEEMS (**Figure 2B**). Further sampling from underrepresented regions—such as Panama, the Gulf side of Mexico, Cuba, and the Windward Islands of the Lesser Antilles—could refine fine-scale migration patterns.

Elsewhere, major oceanographic features also shape gene flow. Expanded sampling in continental Colombia highlights the role of the Colombia-Panama gyre in retaining larvae locally (Baums et al. 2005; García-Urueña et al. 2022). The previous findings of a barrier to gene flow located in the Mona Passage between the Dominican Republic and Puerto Rico is first evident at K = 4 (Baums et al. 2005, 2006b; Galindo et al. 2006). This barrier arises from the combination of oceanographic conditions and life history traits of the species, including the timing of spawning during the summer months and pelagic larval duration (Baums et al. 2006b).

When island habitat is arrayed linearly with respect to major currents, *A. palmata* can exhibit strong isolation-by-distance patterns (**Figure 3**). We consider the analysis presented here exploratory because sampling was not specifically designed to test the hypothesis of isolation-by-distance. However, results are entirely consistent with the findings from other studies in the Antilles (Baums et al. 2005; Devlin-Durante and Baums 2017; Japaud et al. 2019; Alvarado-Cerón et al. 2026). The increase in genetic distance with geographic distance implies that successful reproduction and recruitment is more likely among neighboring sites. It may also indicate that this region is (or was) at an equilibrium between genetic drift (which causes local differentiation) and gene flow (which mixes populations). In any case, the presence of strong isolation-by-distance patterns in this part of the species range makes it challenging to define the location of barriers to gene flow using clustering approaches alone without using additional evidence, e.g. from the biophysical dispersal models mentioned above (Baums et al. 2006b; Ringbauer et al. 2018). Isolation by distance patterns are much weaker along the well-sampled, continuous habitat of the Florida Reef Tract, which may be partially explained by the presence of admixed samples in this region (**Figure 2A**) and partially by the complex current patterns governing local and long-distance dispersal along the reef tract (Dobbelaere et al. 2026).

Although biophysical barriers to gene flow exist, pairwise F_ST_ values indicate overall low differentiation across the Caribbean and north-western Atlantic. They range from 0.001 between Honduras and Belize within the Mesoamerican Reef Tract to 0.125 between US Virgin Islands and Belize/Honduras (**Table 2**). For regions where whole genome and microarray data were available, genome-wide F_ST_ values were similar to microarray-derived F_ST_ values. For example, Florida and Curaçao had a whole genome pairwise F_ST_ value of 0.063 (95% CI [0.061, 0.065], Locatelli et al, unpubl data) and the microarray-derived value here was 0.076 (95% CI [0.071, 0.080]). This low genetic differentiation is in accordance with the natural history of the species. Larvae of *A. palmata* and *A. cervicornis* have a relatively long pre-competency phase and continue to swim upwards in the absence of settlement cues (Limer et al. 2024; Dobbelaere et al. 2026), making long-distance dispersal more likely. Under the assumptions of the Fisher-Wright model, these pairwise F_st_ values translate into between 2 and 250 migrants per generation across sites in the Caribbean and North Western Atlantic **(Table S2)**. The propensity for long-distance dispersal led to a general absence of local kinship **(Figure 4**). Within-site kinship at the second-degree or closer (excluding ramets of the same genet) was only detected at what are presumed to be wild sites in one pair of colonies at Rust Op Twist Reef in the U.S. Virgin Islands, one pair of colonies at Rincon de Guanabo in Cuba, and amongst several colonies at two sites in the Dominican Republic, Catalina and Minitas (all pairwise kinship values corresponded with the expected range for half-siblings or uncle-niece).

The lack of strong population differentiation is underscored by the finding that eggs collected from colonies in Curaçao could be fertilized with cryopreserved sperm from Floridian and Puerto Rican colonies (Hagedorn et al. 2021). This first ever trial of assisted gene flow in a reef-building coral resulted in viable offspring which have been raised for years in closed systems in Florida. A comparison of Curaçao by Florida, Curaçao by Puerto Rico, Florida by Florida and Curaçao by Curaçao colonies revealed that the interpopulation crosses performed at least as well as the Florida by Florida crosses when challenged in a heat stress experiment (Muller et al. 2025).

Assisted gene flow has been identified as a critical intervention tool to prevent genetic diversity loss and support the adaptive potential of endangered coral populations (Baker et al. 2025). Genetic diversity is strongly correlated with species survival and is the most accessible metric for genetic conservation interventions (Reed and Frankham 2003). The interpopulation crosses are incredibly valuable in this respect as they carry a large amount of novel allelic diversity. The Florida x Curaçao colonies harbored over 850 novel alleles that were not present in the Floridian reference population (Muller et al. 2025). This is particularly noteworthy because it has been estimated that *A. palmata* (and *A. cervicornis*) contains roughly 3.5-times lower genome-wide heterozygosity (mean = 0.41% ± 0.02%, Kitchen et al., in prep and Locatelli et al., 2024) compared to other scleractinian corals (mean = 1.42% ± 0.42%; Kitchen et al., in prep). Although these microarray-derived estimates are an order of magnitude higher than genome-wide heterozygosity estimates due to the ascertainment bias of the former (Geibel et al. 2021), our analysis here indicate region-specific variability in heterozygosity (**Table 3**).These findings highlight assisted gene flow as a powerful role tool to enrich genetic diversity via provenancing more distant genets from the species range.

Conservation applications of the presented dataset also include the design of outplanting and breeding strategies. Interventions require decisions on how many and which genets to propagate in *in situ* coral nurseries, over what scale asexually produced fragments and sexually produced larvae may be outplanted (Baums et al. 2019). Because *A. palmata* genets are generally not closely related within sites (**Figure 4**), introduction of sexually produced offspring to the reef of parental origin changes relatedness patterns usually observed in the wild. For example, restoration action has introduced first, second and third degree relatives on a reef in Curacao (Sea Aquarium, Baums et al. 2022). Nursery collections and biobank holdings similarly contain cohorts of recruits that resulted from assisted reproduction (**Figure 4**). Restoration practitioners and managers are well aware of the increased inbreeding risk and are taking steps to track ancestry and optimize outplanting designs to avoid future inbreeding (Margaret M Miller, pers comm). This may include the need to routinely exchange sexually produced offspring between neighboring reefs and islands as well as more distant locations, similar to the natural history of the species.

Caribbean acroporids form large clonal stands through asexual reproduction. Previous analysis of clonal structure across the Caribbean utilized a random, spatially explicit sampling design that standardized sampling effort across sites (Baums et al. 2006a; Japaud et al. 2019). This sampling design revealed a mosaic of sites from those dominated by single genets to those with high genotypic diversity and evenness. Asexual reproduction dominated at the species’ northern range edge, similar to findings in clonal plants and other coral species (Whitaker 2006; Baums et al. 2014). Yet, such systematic analysis of the contribution of asexual reproduction to the population structure of the species is not possible with the current dataset due to a lack of a standardized sampling design. Because coral restoration practitioners are the main users of the tool and often employ STAGdb to confirm the inventory of their coral nurseries, biases in the representation of some genets being represented by up to 75 database records reflect conservation actions rather than the contribution of asexual reproduction to population maintenance in the wild. An updated assessment of clonal structure within the regional and broader populations is needed.

In summary, our comprehensive genome-wide analysis of *A. palmata* reaffirms weak overall genetic differentiation across its range despite the presence of regional structure shaped by biophysical oceanographic features and life history traits. Patterns of admixture, long-distance dispersal, and low kinship underscore the high connectivity among the nine populations. These findings highlight both the resilience and vulnerability of the species.

Although gene flow was sufficiently high to maintain connectivity in the past, continued population declines threaten the preservation of genetic diversity. Importantly, our results refine conservation strategies by supporting assisted gene flow as a tool to enhance genetic diversity and avoid potential future inbreeding as a consequence of restoration interventions. In the future, we recommend integrating standardized sampling designs with genomic monitoring to guide effective, evidence-based restoration and management efforts across the Caribbean.

## Author contributions

IBB conceived the study, interpreted results and wrote the manuscript. NSL, KLdL and SAK performed data analysis and interpretation. All authors edited the manuscript.

## Acknowledgements

We gratefully acknowledge the STAGdb contributors who have generously made their data publicly available, in particular individuals at the following organizations: Centro de Investigaciones Marinas (CIM-UH); Coral Morphologic; Coral Restoration Foundation; Florida Aquarium (FLAQ); Florida Fish and Wildlife Conservation Commission (FWC); Iberostar Wave of Change; Mote Marine Laboratory; National Oceanic and Atmospheric Administration (NOAA); Nova Southeastern University; National Park Services (NPS); Oceanus; Pennsylvania State University; Rare; Reef Renewal USA; Reef Renewal Foundation Curacao; Sea Ventures; SECORE International; The Nature Conservancy; The Reef Institute; Universidad Nacional Autónoma de México (UNAM); University of Magdalena; University of Miami; Washington and Lee University; and any others that we were unable to identify by name. We further thank Matthias Schröder (UOL ICBM) for technical support.

## Data availability

Coral genotype calls and associated metadata are available through the STAGdb Galaxy resource at coralsnp.uol.de..

## Supplements

### Supplemental Figures

**Figure S1.**
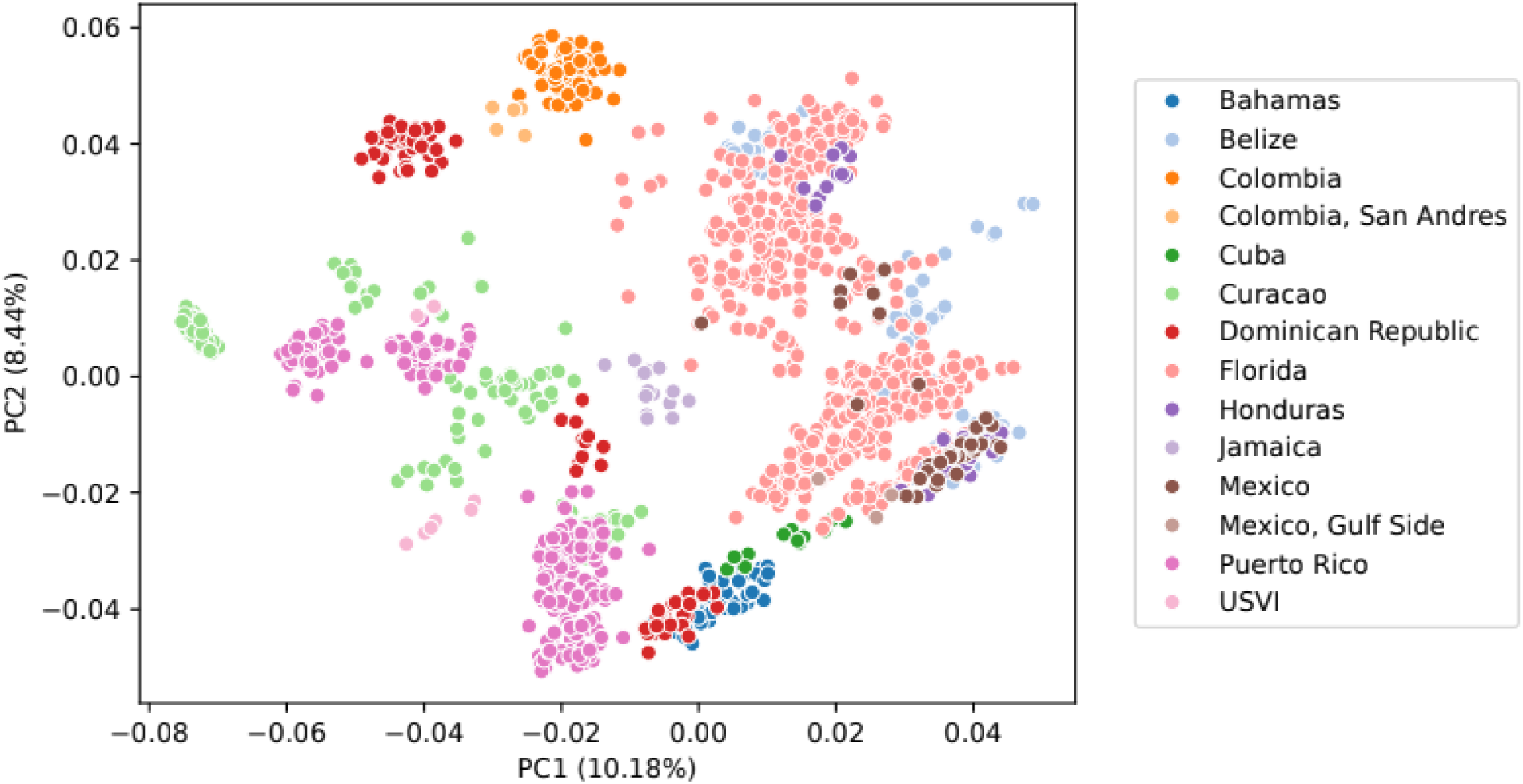
Principal component analysis of STAGdb data from *Acropora palmata* genets using all genotyping probes instead of only the best and recommended probe set (minor allele frequency threshold of 0.05, dataset 5).

**Figure S2.**
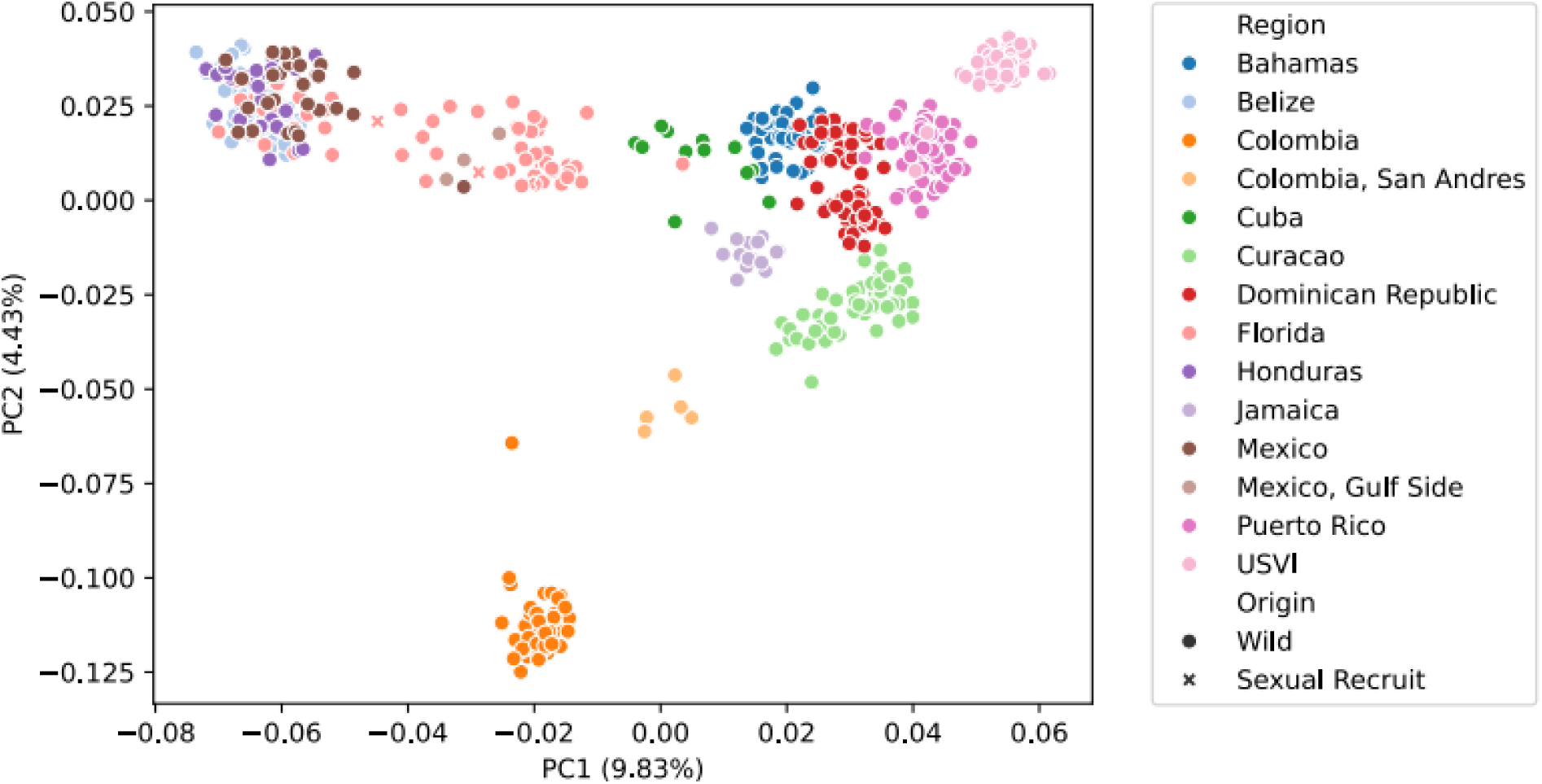
Principal component analysis of STAGdb data from *Acropora palmata* genets without removing low confidence SNP calls. Note the split of samples from the Dominican Republic into two clusters that are not present after removing low confidence SNP calls (dataset 6, Fig 1). K=2 (optimal by deltaK):

**Figure S3.**
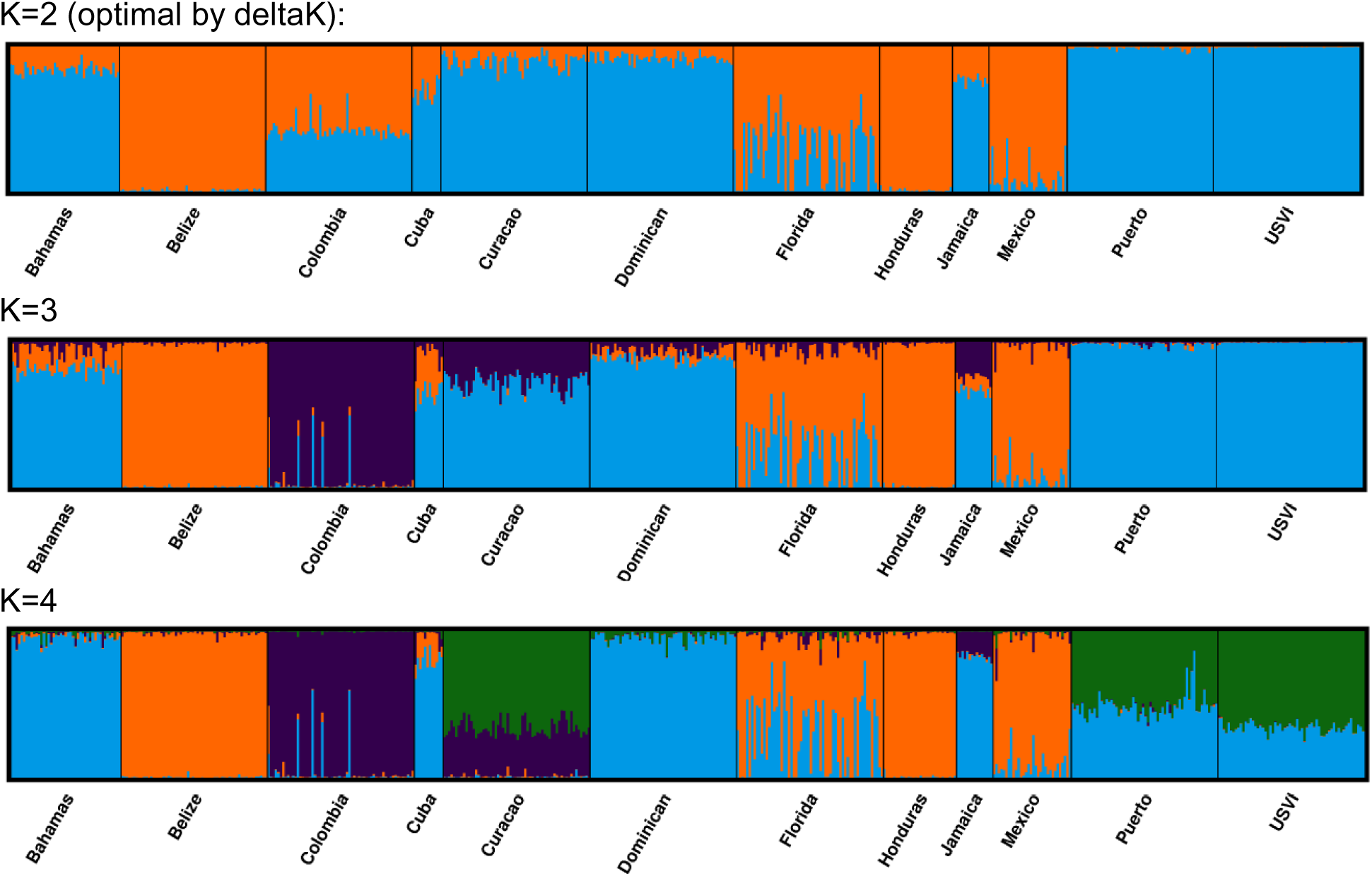
STRUCTURE output from K=2 to K=4 showing the hierarchical population structure of *A. palmata*. The optimal K according to delta K method is K=2. K=7 (CV error optimal) ADMIXTURE K=9 (Puechmaille optimal) ADMIXTURE

**Figure S4.**
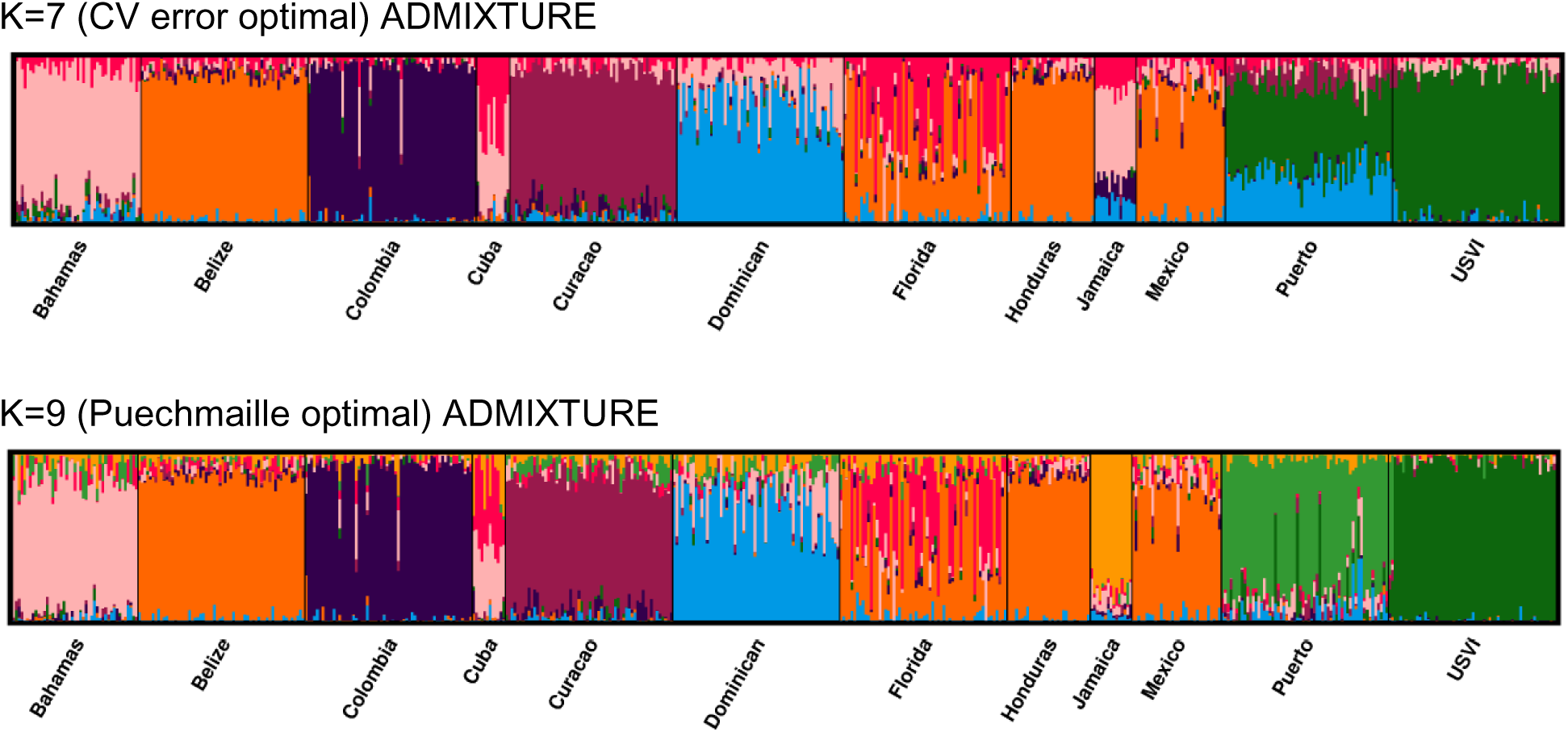
ADMIXTURE plots for K=7 and K=9. The optimal K values differed according to metric tested, K=7 was the optimal according to the CV error method and K=9 was the optimal K according to the Puechmaille method.

### Tables

**Table S1. Metadata table including the columns:** [affymetrix_id, species, mlg, region, region_sub, region_fix_reason, reef, lat_old, lon_old, lat_clean, lon_clean, coord_fix_reason, is_nursery, filtered_genets_n554, sexual_recruit, sample_id, user_specimen_id, perc_missing_data, perc_heterozygous, perc_acer, perc_apal, seq_facility, organization, collector_last_name, collection_date]

**Table S2.**
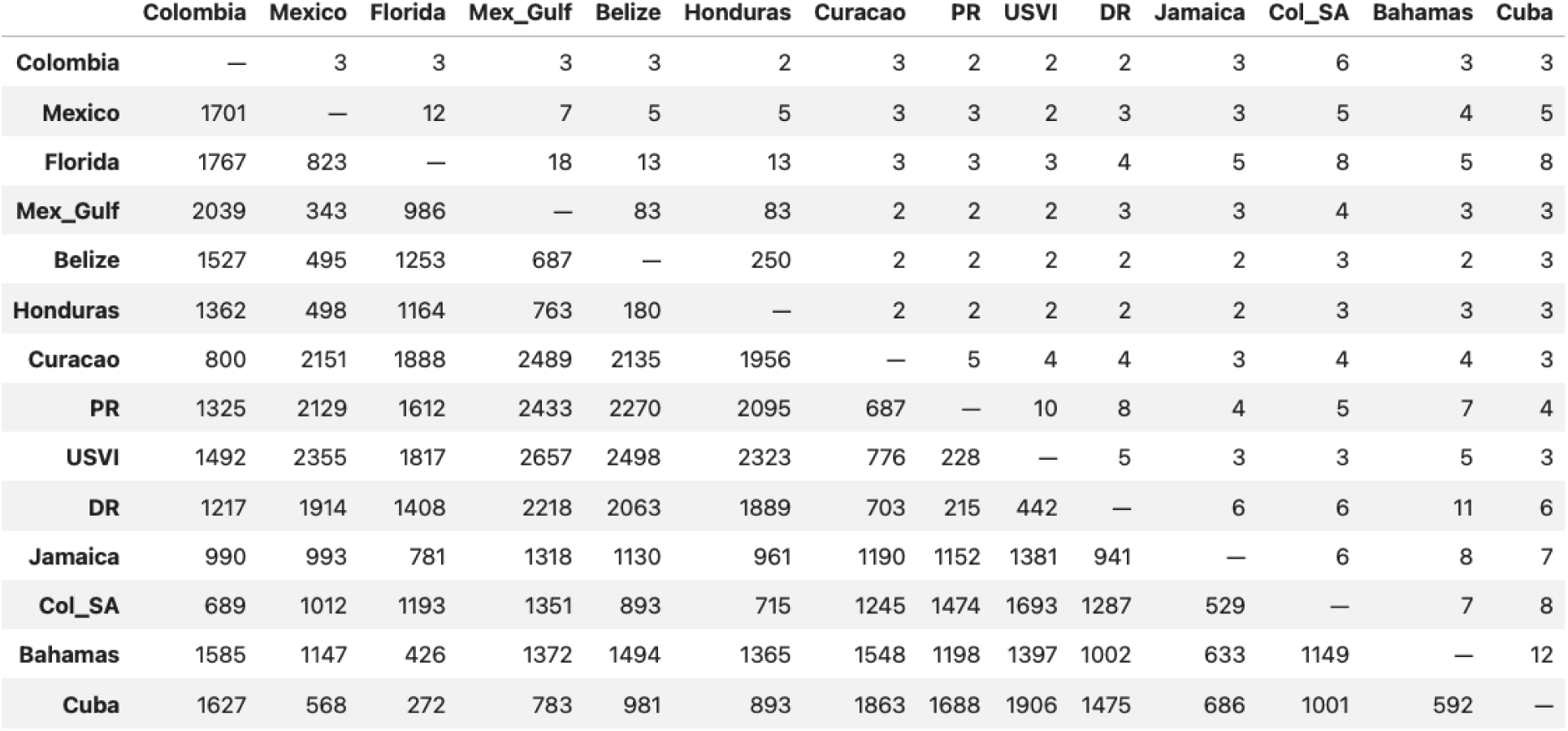
Pairwise estimates of migrants (upper triangle) and geographic distance (lower triangle). The number of migrants is derived from pairwise F_ST_ calculated under Wright’s island model. Rows and columns are ordered by hierarchical clustering of the pairwise F_ST_ matrix to group genetically similar regions. Values should be interpreted as a gene flow proxy rather than as direct estimates of realized migration.

|  | Colombia | Mexico | Florida | Mex_Gulf | Belize | Honduras | Curacao | PR | USVI | DR | Jamaica | Col_SA | Bahamas | Cuba |
| --- | --- | --- | --- | --- | --- | --- | --- | --- | --- | --- | --- | --- | --- | --- |
| Colombia | — | 3 | 3 | 3 | 3 | 2 | 3 | 2 | 2 | 2 | 3 | 6 | 3 | 3 |
| Mexico | 1701 | — | 12 | 7 | 5 | 5 | 3 | 3 | 2 | 3 | 3 | 5 | 4 | 5 |
| Florida | 1767 | 823 | — | 18 | 13 | 13 | 3 | 3 | 3 | 4 | 5 | 8 | 5 | 8 |
| Mex_Gulf | 2039 | 343 | 986 | — | 83 | 83 | 2 | 2 | 2 | 3 | 3 | 4 | 3 | 3 |
| Belize | 1527 | 495 | 1253 | 687 | — | 250 | 2 | 2 | 2 | 2 | 2 | 3 | 2 | 3 |
| Honduras | 1362 | 498 | 1164 | 763 | 180 | — | 2 | 2 | 2 | 2 | 2 | 3 | 3 | 3 |
| Curacao | 800 | 2151 | 1888 | 2489 | 2135 | 1956 | — | 5 | 4 | 4 | 3 | 4 | 4 | 3 |
| PR | 1325 | 2129 | 1612 | 2433 | 2270 | 2095 | 687 | — | 10 | 8 | 4 | 5 | 7 | 4 |
| USVI | 1492 | 2355 | 1817 | 2657 | 2498 | 2323 | 776 | 228 | — | 5 | 3 | 3 | 5 | 3 |
| DR | 1217 | 1914 | 1408 | 2218 | 2063 | 1889 | 703 | 215 | 442 | — | 6 | 6 | 11 | 6 |
| Jamaica | 990 | 993 | 781 | 1318 | 1130 | 961 | 1190 | 1152 | 1381 | 941 | — | 6 | 8 | 7 |
| Col_SA | 689 | 1012 | 1193 | 1351 | 893 | 715 | 1245 | 1474 | 1693 | 1287 | 529 | — | 7 | 8 |
| Bahamas | 1585 | 1147 | 426 | 1372 | 1494 | 1365 | 1548 | 1198 | 1397 | 1002 | 633 | 1149 | — | 12 |
| Cuba | 1627 | 568 | 272 | 783 | 981 | 893 | 1863 | 1688 | 1906 | 1475 | 686 | 1001 | 592 | — |

